# Inbred or Outbred? Genetic diversity in laboratory rodent colonies

**DOI:** 10.1101/174102

**Authors:** Thomas D. Brekke, Katherine A. Steele, John F. Mulley

## Abstract

Non-model rodents are widely used as subjects for both basic and applied biological research, but the genetic diversity of the study individuals is rarely quantified. University-housed colonies tend to be small and subject to founder effects and genetic drift and so may be highly inbred or show substantial genetic divergence from other colonies, even those derived from the same source. Disregard for the levels of genetic diversity in an animal colony may result in a failure to replicate results if a different colony is used to repeat an experiment, as different colonies may have fixed alternative variants. Here we use high throughput sequencing to demonstrate genetic divergence in three isolated colonies of Mongolian gerbil (*Meriones unguiculatus*) even though they were all established recently from the same source. We also show that genetic diversity in allegedly ‘outbred’ colonies of non-model rodents (gerbils, hamsters, house mice, and deer mice) varies considerably from nearly no segregating diversity, to very high levels of polymorphism. We conclude that genetic divergence in isolated colonies may play an important role in the ‘replication crisis’. In a more positive light, divergent rodent colonies represent an opportunity to leverage genetically distinct individuals in genetic crossing experiments. In sum, awareness of the genetic diversity of an animal colony is paramount as it allows researchers to properly replicate experiments and also to capitalize on other, genetically distinct individuals to explore the genetic basis of a trait.

## INTRODUCTION

The genetic variation present in laboratory rodent colonies has important implications for the design, outcome and reproducibility of biological experiments (Justice & Dhillon, 2016). High levels of genetic variation reduce power and increase variation in the response to a treatment, but the experimental results may be more applicable to natural or human populations. Alternatively, inbred colonies provide more power and require fewer animals per experiment by limiting the noise caused by segregating genetic variation (here we define an inbred strain as the result of ≥20 generations of brother-sister mating or equivalent (Casellas, 2011; Eppig, 2007)). Indeed, minimizing the number of animals (in accordance with the principle of *reduction* in the 3Rs (Russell & Burch, 1959)) is one of the main reasons cited for the use of inbred lines rather than outbred colonies (Chia, Achilli, Festing, & Fisher, 2005; Festing, 1999; Groen & Lagerwerf, 1979). Inbred lines with single genes knocked out have proven tremendously powerful for identifying the phenotypic effect of those genes (Festing, 2010), but phenotypic traits and diseases often have complex genetic bases (e.g.: diabetes (Fuchsberger et al., 2016; Rich, 2016), and epilepsy (Meisler, Kearney, Ottman, & Escayg, 2001)), so inbred models with no genetic variation may preclude a complete understanding of the underlying genetic architecture. Genetic variation is essential for the identification of candidate genes underlying complex phenotypes, and projects such as the collaborative cross have gone to great effort (and expense) to reestablish segregating variation into inbred mouse strains in a controlled manner (Churchill et al., 2004; Collaborative Cross Consortium, 2012; Threadgill & Churchill, 2012). Such projects rely on the fact that while there is no segregating variation within a single inbred line, multiple inbred lines have fixed alternative variants and immense power can be gained by leveraging these fixed alleles in a genetic mapping experiment (Collaborative Cross Consortium, 2012; de Koning & McIntyre, 2017; Svenson, Gatti, Valdar, & Welsh, 2012).

Genetic crosses involving multiple inbred lines are hugely powerful for genetic experiments, but true inbred strains of mammals are rare outside of ‘model’ rodents such as mice and rats. The use of ‘non-model’ rodents, such as gerbils (*Meriones unguiculatus*, (Stuermer & Wetzel, 2006), hamsters (*Phodopus sp.*, (Brekke, Henry, & Good, 2016), spiny mice (*Acomys sp.*, (Gawriluk et al., 2016) and deer mice (*Peromyscus sp.*, (Weber, Peterson, & Hoekstra, 2013), is mainly restricted to outbred colonies with standing genetic variation. Unfortunately, even in outbred strains of house mice genetic diversity is often ill-defined (Chia et al., 2005), and surprisingly little work has been done to quantify diversity in colonies of non-model rodents. Indeed, the labeling of a strain of animals as ‘outbred’ (Chia et al., 2005) or ‘wild-derived’ (Harper, 2008) may have little to no bearing on the genetic diversity present. Instead, such labels only demonstrate that the animals have not *purposely* undergone the ≥20 generations of brother-sister mating necessary to purge segregating variation and establish a true inbred line (Casellas, 2011; Eppig, 2007). Furthermore, while it is recognized that large colonies will slow the loss of genetic variation through drift (Papaioannou & Festing, 1980), and commercial providers of outbred animals may maintain 50-100 breeding pairs per colony, the size of colonies in academic institutions is constrained by housing space, finances, and human resources. Furthermore, bottleneck or founder effects are likely to occur as animals are moved between colonies, or used to establish a new one. Thus, we should expect the amount of standing genetic variation to differ even between colonies of the same species and strain.

Mongolian gerbils (*Meriones unguiculatus*) are a common non-model rodent that have been used in biological research for many years and have informed our understanding of diseases such as epilepsy (Buchhalter, 1993; Buckmaster, 2006), stroke (Vincent & Rodrick, 1979), and diabetes (X. Li et al., 2016) as well as basic biology questions about thermal regulation (Thiessen & Kittrell, 1980; D. Wang, Wang, & Wang, 2000), desert adaptation (McManus, 1972), domestication (Stuermer & Wetzel, 2006; Stuermer et al., 2003), reproductive biology (Clark, Ham, & Galef, 1994), hearing (Abbas & Rivolta, 2015; Chen et al., 2012), and more. Despite the widespread use of gerbils in scientific research, few widely-accessible transcriptomic and genomic resources have been developed, and the small numbers of genetic markers available are severely limited in their ability to reveal levels of genetic diversity in laboratory gerbils. Early reports using microsatellites (Neumann, Maak, & Stuermer, 2001) and AFLPs (Razzoli, Papa, Valsecchi, & Nonnis Marzano, 2003) suggested that genetic diversity in laboratory gerbil colonies is a small fraction of that in the wild, and below that of inbred mouse or rat strains, but more recent reports, also using microsatellites, suggest that variation is quite high (Du et al., 2015; 2010). A simple explanation for this contradiction is that different strains of animals were used in each study. Du et al. (2010 and 2015) surveyed four laboratory colonies from China, all of which were recently established from wild caught individuals, whilst Neumann et al. (2001) used animals originating from the Tumblebrook Farm strain. This strain has its origins in 20 pairs of wild-caught animals that were used to establish a colony in the Kitasato Institute in Japan in 1935. An unknown number of gerbils were subsequently transferred to the Central Laboratories for Experimental Animals in 1949 (Petrij, van Veen, Mettler, & Brückmann, 2001). In 1954, eleven pairs of these animals were transferred to Tumblebrook Farm in the USA (of which five females and four males reproduced (Stuermer et al., 2003)), forming the basis of what might be thought of as the “domesticated laboratory gerbil” (*Meriones unguiculatus forma domestica*,(Stuermer et al., 2003)). Later the Tumblebrook colony was purchased by Charles River Ltd in 1996 and the strain was then rederived and maintained in Italy (Neumann et al., 2001; Razzoli et al., 2003). These Tumblebrook animals have been maintained since then as an outbred colony with ≥100 breeding pairs (C. Parady, personal communication). The population history of laboratory gerbils is punctuated by a series of bottleneck events each time the colony was moved and rederived. There is therefore a discrepancy in how this animal is maintained and sold by commercial providers (as a highly diverse outbred stock) and the results of previous genetic analyses (which suggest very low levels of diversity (Neumann et al., 2001; Razzoli et al., 2003)). If gerbils are inbred, fewer are needed to achieve statistically significant results and maintain a breeding colony. Given the limitations of small-scale microsatellite and AFLP experiments, we therefore decided to use a genome-wide approach to quantify the genetic diversity present in Tumblebrook Farm strain gerbils. Here, we evaluate patterns of standing genetic variation in animals from three different gerbil colonies to identify differences that may stem from a history of bottlenecks and isolation. All three colonies originated from the European colony managed by Charles River Ltd. We also compared these with the recently released whole genome sequence of an individual from an American stock of the Tumblebrook Farm strain (genbank accession GCA_002204375.1). We interpret the levels of genetic variation in gerbils in comparison with colonies of other species such as house mice (*Mus musculus ssp.*), hamsters (*Phodopus sp*.), and deer mice (*Peromyscus sp*.). We also discuss the possibility of leveraging the genetic drift inescapable in small mammal colonies to identify differentiated genetic markers for use in genetic mapping and association studies.

## MATERIALS AND METHODS

### Animals

Mongolian gerbils are listed in Annex 1 of EU Directive 2010/63/EU and must therefore be purposely bred for scientific research. The majority (if not all) gerbils used in the European Union are derived from the Tumblebrook farm stock and many academic institutions in the UK and elsewhere maintain their own colonies derived from these animals. We analyzed animals from three of these colonies, and to avoid confusion we refer to each colony by the name of the city where it was first established: Edinburgh, Sheffield, and Bangor. The Edinburgh colony was established by Dr. Judith Allen at the University of Edinburgh circa 2005. In 2014, Dr. Leila Abbas established a new colony at the University of Sheffield from the Charles River Ltd Tumblebrook stock and at the same time took over care and housing of 3-4 pairs of animals from Edinburgh, with both stocks maintained separately. In 2016, we took delivery of 12 new Tumblebrook animals from Charles River (7♀, 5♂) to establish the Bangor colony. We also received 5 animals from each of the Edinburgh (3♀, 2♂) and Sheffield (2♀, 3♂) stocks. All three groups were maintained in isolation in Bangor, except for a single test cross between an Edinburgh female and a Sheffield male. All animals were housed in accordance with E.U. and Home Office animal care regulations and experiments were reviewed and approved by the Bangor University Animal Welfare and Ethical Review Board.

### Tissue collection, DNA extraction, library preparation, and sequencing

Liver tissue was collected from the 22 founder animals and 2 F_1_ Edinburgh x Sheffield offspring and snap-frozen immediately in liquid nitrogen after each animal was euthanized as part of routine colony management. DNA was extracted with the Qiagen DNAeasy Blood & Tissue kit and treated with RNase according to manufacturer’s instructions. Extracted DNA was shipped BGI (Hong Kong) for library preparation and sequencing. Uniquely barcoded 100 base-pair paired-end genotyping-by-sequencing (GBS) libraries were prepared with the 5 base-pair cutter ApeKI, pooled, and sequenced on a single lane of Illumina 4000 (Elshire et al., 2011).

### Bioinformatics

BGI filtered the raw data through their SOAPnuke filter, which includes demultiplexing the reads and removing proprietary barcode sequences, and dropping reads that were >26% adapter sequence, and/or >40% of the bases below a PHRED quality score of 15. We used the Stacks (v1.46 Catchen, Amores, Hohenlohe, Cresko, & Postlethwait, 2011; Catchen, Hohenlohe, Bassham, Amores, & Cresko, 2013) pipeline to identify tags and call SNPs from the first reads, resulting in an average sequencing effort of 7.8 million reads per individual. This analysis included the standard Stacks pipeline components: process_radtags, ustacks, cstacks, sstacks, and populations. All scripts were run with default flags with the following exceptions. With process_radtags we cleaned reads (-c), discarded reads with low quality scores (-q), rescued radtags (-r), and truncated read length to 92 bases (-t 92) to avoid variation in read length that would otherwise disrupt the remaining pipeline. Thus, our final markers were all 92 base-pairs long. We ran the deleveraging algorithm in ustacks (-d) and used 6 individuals (a female and male from each Bangor, Edinburgh, and Sheffield strains) for cstacks. We generated a reference fasta from the output of cstacks and to it we aligned the raw reads with bwa mem (H. Li & Durbin, 2009). From these alignments, we extracted depth of coverage with samtools (H. Li et al., 2009) in order to annotate autosomal, X-, and Y-linked markers. Coverage was standardized by the sequencing effort of each individual and multiplied by 1,000,000 before being summed across males and females. Sex-linkage is apparent by comparing standardized coverage of each marker in males versus females. We first annotated markers with less than 10x total standardized coverage as ‘unknown’ and removed from the dataset as these have too low coverage to reliably differentiate X- and Y-linked tags from autosomal tags or call variants. We next identified Y-linked markers as those with <1x standardized coverage in females. X-linked markers fulfilled the inequality: Coverage^male^ < ¾ Coverage^female^ − 5. The slope of this line was chosen to discriminate points in the X-linked cluster (slope = ½) from those in the autosomal cluster (slope = 1). The intercept was chosen in order to remain fairly conservative near the origin; that is erroring towards labeling a true X-linked tag as an autosome rather than labeling a true autosomal tag as X-linked. All remaining tags were annotated as autosomal. The populations script was used to generate diversity metrics and F statistics across the genome for autosomal, X-, and Y-linked markers. Genotypes were called only for individuals with greater than 10x coverage (-m 10) and only for SNPs in the first 90 bases. We blacklisted SNPs in the final two bases because of an unusually high number of SNPs on those bases (for further discussion see Supplemental Figure 1). Finally, we evaluated genetic similarity and population structure with the SNPRelate package in R (Zheng, Levine, Shen, & Gogarten, 2012) and the program Structure (Falush, Stephens, & Pritchard, 2003; 2007; Pritchard, Stephens, & Donnelly, 2000; Rosenberg et al., 2002). We visualized structure data with the program distruct (Rosenberg, 2003). We calculated pairwise diversity between our reference and the recently released gerbil whole genome sequence (genbank accession GCA_002204375.1) by aligning the reference sequences to the genome with bwa mem, discarding partial-length alignments and counting mismatches across the first 90 bases of the reference.

In order to evaluate the levels of nucleotide diversity in gerbils in a more general sense, we downloaded RAD sequencing data from deer mice (SRA accession PRJNA186607; Weber et al., 2013) and were provided RAD sequence data for hamsters (Jeff Good, personal communication). These RAD datasets were analyzed with the Stacks pipeline described above (omitting the chromosomal annotation steps) in order to be directly comparable with diversity estimates in gerbils. To provide further context for our estimates of π we also retrieved recently published diversity metrics from house mice in the collaborative cross (Srivastava et al., 2017).

### Data availability

Raw sequencing data is archived in the SRA under the BioProject accession number PRJNA397533 and the sample accession numbers SAMN07460176-SAMN07460199. The reference fasta file (including Autosomal, X-, or Y-linkage of each tag) as well as the VCF file containing the locations of all SNPs are available as supplemental data.

## RESULTS

We compared sequencing coverage in females and males to annotate 718,385 autosomal markers, 5,148 X-linked markers, and 2,355 Y-linked markers (Figure 1). We identified 30,365 SNPs spread across 24,326 markers (1.25 SNPs/marker). Average autosomal nucleotide diversity (π) is 0.0059 (Table 1), which describes the variation in unconstrained, non-coding regions. Average heterozygosity at autosomal variant sites is 0.447 and is slightly higher on the sex chromosomes (Table 1).

**Figure 1.**
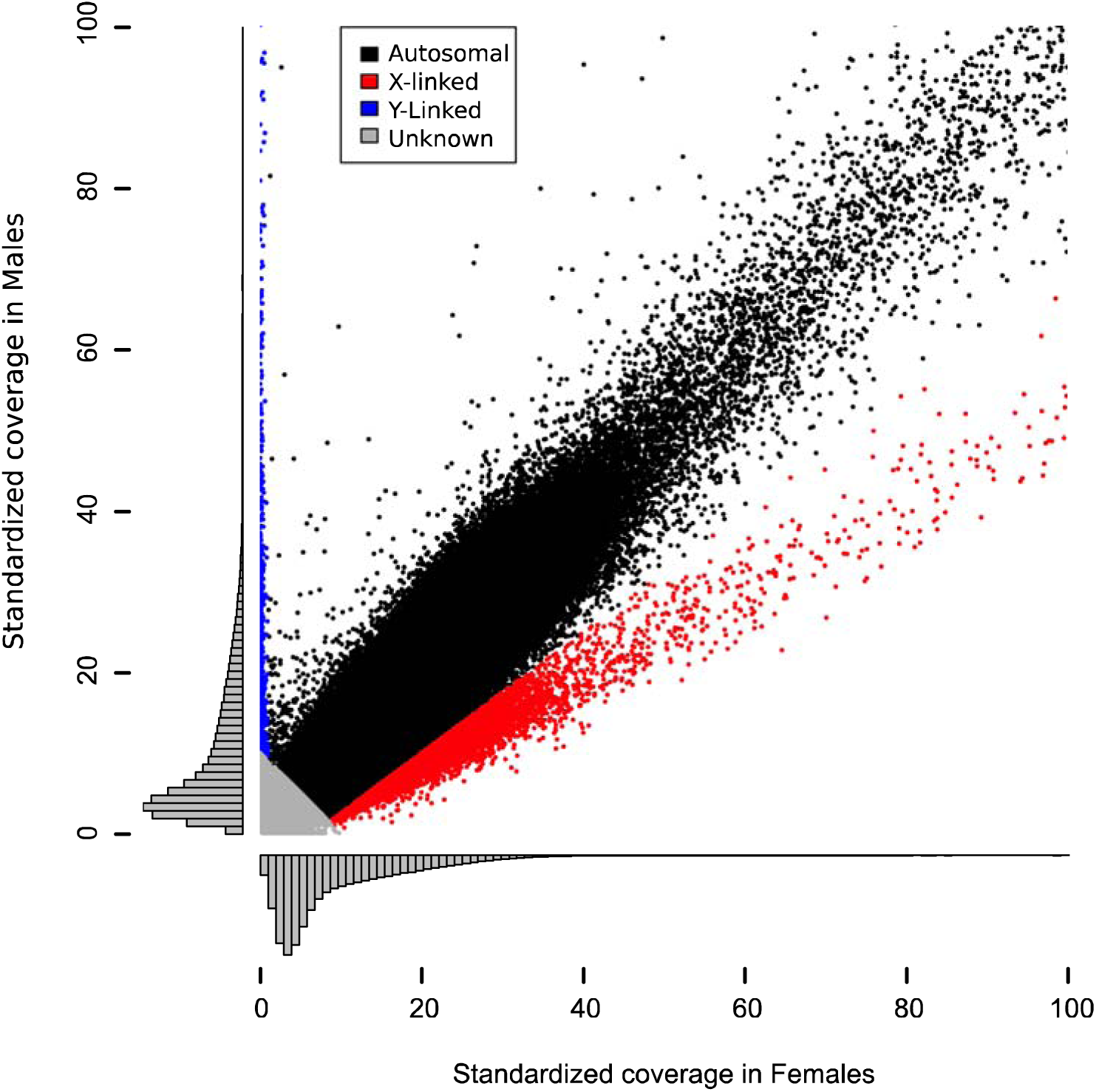
Relative coverage in females and males can be used to identify sex chromosomes. Plotted here is standardized coverage in females against standardized coverage in males, histograms show the density of markers along each axis. The markers shown are those with low overall coverage; a long tail exists in both females and males and is not shown here. markers with less than 10x total standardized coverage were annotated as unknown (grey) because those have too little coverage to reliably distinguish X- and Y-linkage from autosomes. Of the remaining tags, those with less than 1x standardized coverage in females are annotated as Y-linked (blue). Those which satisfy the inequality: Coverage^male^ < ¾ Coverage^female^ − 5 were identified as X-linked (red). This line has a slope designed to discriminate points in the X-linked cluster (slope = ½) from those in the autosomal cluster (slope = 1) while remaining fairly conservative near the origin. All remaining tags were annotated as autosomal (black).

**Table 1.**
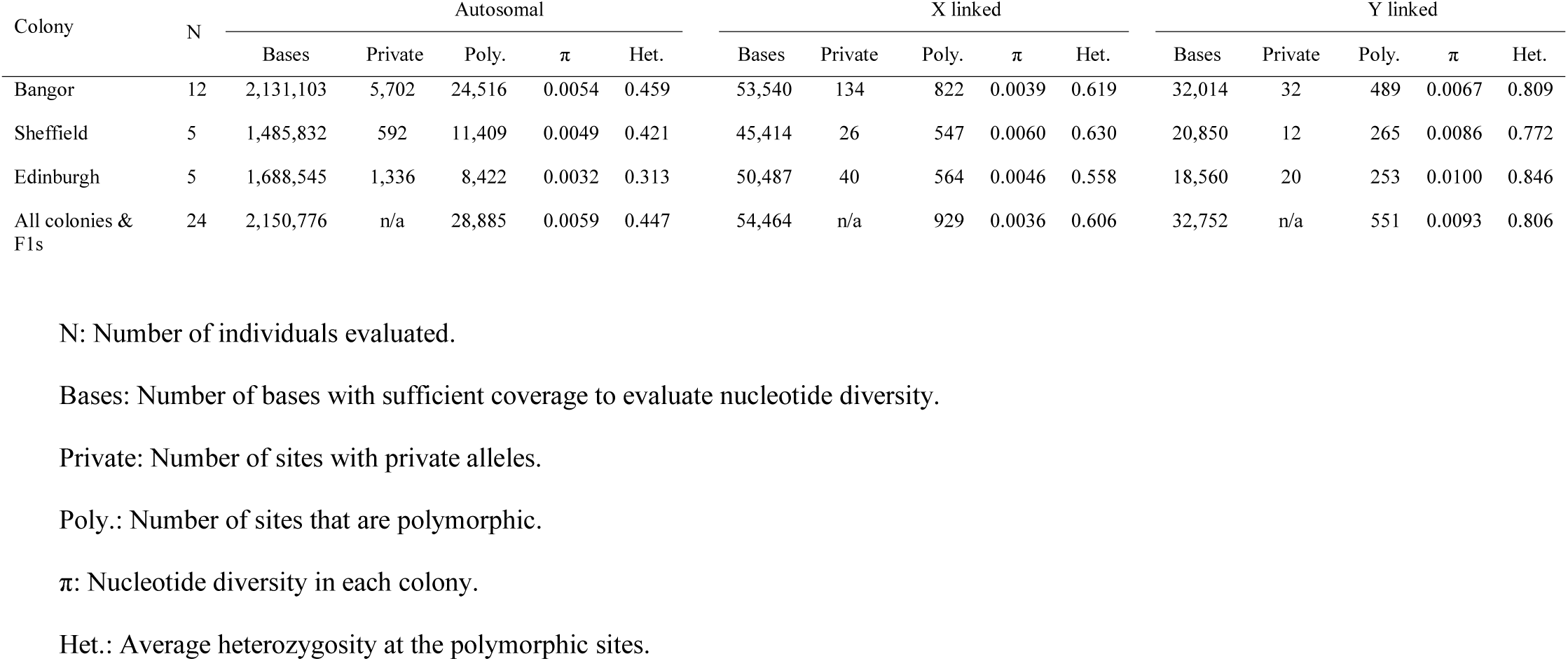
Diversity metrics in gerbils across the genome.

In order to evaluate how different the gerbil genome (GCA_002204375.1) is from our colonies, we counted the number of differences between our GBS reference and the genome. Full-length alignments were found for 674,342 of our reference sequences when aligned to the genome. We found 47,223 single-base differences in these aligned regions, far more SNPs than segregate within the colonies we assayed. This pattern is consistent with the known population history of laboratory gerbils: the Charles River colony, from which our animals originate, was rederived from a U.S. colony from which the DNA for the genome was supplied.

We used 28,885 autosomal SNPs to evaluate the diversity between the three colonies and found that while each colony does possess a small set of private alleles, most alleles are shared across all colonies (Table 1). A substantial portion of the variation (24%) is explained by differences between Edinburgh and the other colonies (Figure 2). Eigenvector 2 shows that much of the remaining variation (7%) segregates within the Bangor colony. No higher-order eigenvectors discriminate the colonies, instead they partition variation common to all. F_st_ metrics between the colonies suggest high overall similarity between Bangor and Sheffield (F_st_ = 0.069) while identifying Edinburgh as an outlier with F_st_ of 0.235 compared to Bangor and 0.352 to Sheffield. The structure analysis also suggests little overall differentiation between Bangor and Sheffield, and finds that Edinburgh is slightly more divergent, though still very similar (Figure 3). Overall, these data suggest that while Edinburgh animals have marked differences from other gerbils, they still share many genetic variants.

**Figure 2.**
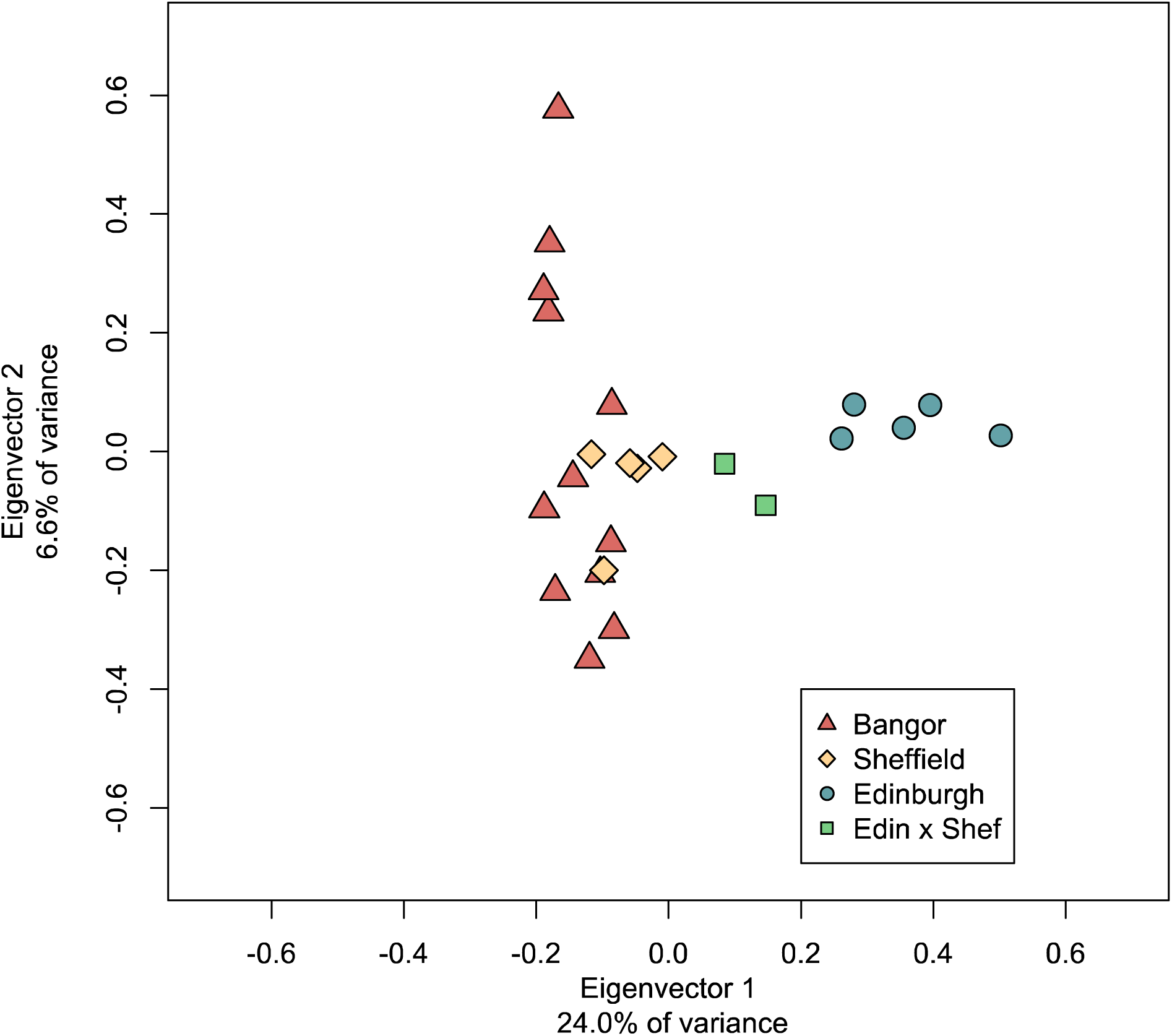
PCA of 28,885 autosomal SNPs. Eigenvector 1 explains the majority of the diversity in these samples and strongly differentiates the Edinburgh colony from the others. Sheffield contains a subset of the genetic diversity found within Bangor colony. F1 offspring between Edinburgh female and a Sheffield male fall out intermediate.

**Figure 3.**
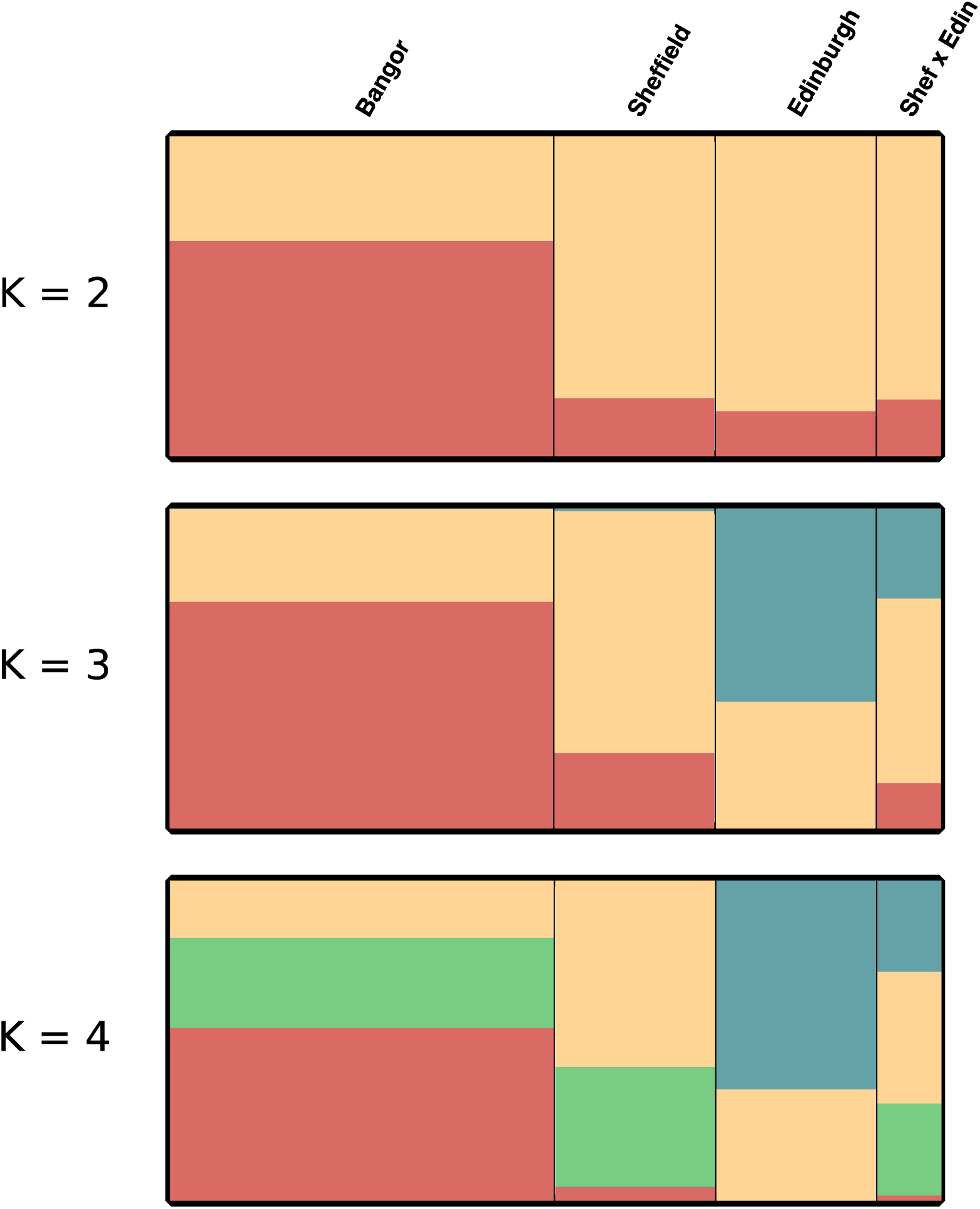
Structure plots of gerbil colonies. Structure was run for 20,000 iterations with 10,000 iterations of burn-in for K=2, 3, and 4. In general, the different colonies have similar assignment especially at low K values. Edinburgh animals fall out uniquely at higher K values (i.e.: blue). The F1 hybrids have mixed ancestry as expected for the offspring of a Sheffield x Edinburgh cross.

In general, the highest diversity is found in the Bangor animals. This is apparent in both the PCA (Figure 2) and the number of polymorphic sites segregating within the Bangor strain (Table 1) and may be due to the higher number of individuals screened. Edinburgh animals have far fewer segregating sites, lower nucleotide diversity, and lower heterozygosity overall, which may be a result of serial bottlenecks. The principle components analysis and the low number of private alleles suggests that the Sheffield animals contain a subset of the diversity found within the Bangor strain (Figure 2). As expected the F1 offspring between Edinburgh and Sheffield are found intermediate to the parents.

Nucleotide diversity in gerbils is quite high compared with other laboratory rodents (Table 2). In fact, genetic diversity in the gerbils often rivals the diversity found in wild-caught mice, and, contrary to previous claims (Neumann et al., 2001; Razzoli et al., 2003), far exceeds the diversity in inbred mice and rats (Ness et al., 2012; Salcedo, Geraldes, & Nachman, 2007; Smits, van Zutphen, Plasterk, & Cuppen, 2004). It is clear that genetic diversity in Tumblebrook gerbils is much higher than previous reports suggest (Neumann et al., 2001; Razzoli et al., 2003). It is also clear that the breeding scheme alone does not robustly predict the amount of standing genetic diversity of an animal colony, especially in non-model rodents.

**Table 2.** Nucleotide diversity in various rodent colonies.

| Species | $\pi$ | Number of individuals | Breeding scheme | Region of genome for which $\pi$ was evaluated | Approximate number of bases surveyed | Citation |
| --- | --- | --- | --- | --- | --- | --- |
| *Meriones unguiculatus* | 0.0059 | 24 | Outbred | Near autosomal restriction sites (GBS) | 2,200,000 | This paper |
| *Mus musculus ssp.*<sup>a</sup> | 0.0289 | 69 | Collaborative cross | Whole genome sequencing | 2,300,000,000 | (Srivastava et al., 2017) |
| *Peromyscus maniculatus*<sup>b</sup> | 0.0006 | 13 | Outbred | Near restriction sites (ddRAD) | 400,000 | (Weber et al., 2013) |
| *Peromyscus polionotus*<sup>b</sup> | 0.0010 | 1 | Outbred | Near restriction sites (ddRAD) | 400,000 | (Weber et al., 2013) |
| *Phodopus campbelli*<sup>c</sup> | 0.0006 | 14 | Outbred | Near restriction sites (ddRAD) | 1,300,000 | J. Good, personal communication |
| *Phodopus sungorus*<sup>c</sup> | 0.0002 | 11 | Outbred | Near restriction sites (ddRAD) | 1,400,000 | J. Good, personal communication |

<sup>a</sup> Based on mean value of 69 collaborative cross individuals. Data from the column titled “% het (autosomes) in sequenced sample” in Supplemental Table 2 of (Srivastava et al., 2017). Approximate number of bases surveyed is based on a genome size of 2.7 Gigabases divided by the mean “% coverage at 15x”.
<sup>b</sup> The *Peromyscus* animals evaluated here are the BW and PO strains originating with the *Peromyscus* stock center, bred at Harvard University in the Hoekstra lab, and sequenced by (Weber et al., 2013).
<sup>c</sup> The *Phodopus* animals evaluated here are from the Good Lab at the University of Montana described in (Brekke & Good, 2014).

## DISCUSSION

We have demonstrated that appreciable genetic diversity segregates within Tumblebrook Farm strain Mongolian gerbils. Our findings are contrary to earlier reports suggesting that diversity may be as low as, or lower than inbred mouse colonies (Neumann et al., 2001; Razzoli et al., 2003). These reports evaluated a small portion of the genome and, perhaps unsurprisingly, found little variation. For instance, Neumann et al. (2001) evaluated diversity at nine microsatellites and found that lab strains had severely reduced allelic diversity compared to wild animals and Razzoli et al. (2003) found a similar pattern using 228 AFLP fragments from six primer combinations. Our genome-wide assay evaluated millions of bases and so has much higher power to find rare variants. Using these data we find that genetic diversity in Mongolian gerbils is relatively high amongst outbred rodent colonies (π = 0.0059 in gerbils and π ≤ 0.0010 in other rodents, Table 2).

Lab-maintained rodent colonies are often small due to the costs and space needed for maintenance of many animals. With such small populations, genetic drift plays an important role in determining the standing level of variation. Drift can be expected to increase genetic differentiation between colonies through time, especially given the population bottleneck that often occurs when a colony is established or moved to a new location. Knowledge of levels of genetic diversity in an institutional colony is therefore vital for correct colony management – for example *Phodopus* hamsters have been referred to as outbred and maintained in large colonies (Brekke & Good, 2014). However, analysis of ddRAD data from two hamster species (J. Good, personal communication) shows that in fact genetic diversity is extremely low in both (Table 2), and so hamster colonies could be maintained with few individuals with no resulting loss of diversity. Despite the length of time in captivity, the Tumblebrook gerbils are (correctly) maintained as a large outbred colony (≥100 breeding pairs) by Charles River Ltd. Our data suggest that the diversity present in that original stock has been sub-sampled and exposed to drift in each of the three independent colonies we assayed. At one extreme is the Edinburgh colony which was not only the first isolated from Tumblebrook, but has been transferred through three universities and experienced the associated bottlenecks. Given this history, it is not surprising that the Edinburgh animals have the fewest SNPs segregating within them, nor that they are somewhat differentiated from Bangor and Sheffield. The Sheffield animals, which were established from Tumblebrook strain founders in 2014 and have been moved through only two universities, also show a reduced diversity, though still higher than Edinburgh. The Bangor colony was established most recently in 2016 and has the highest amount of diversity. As these animals were sent directly from Charles River Ltd, they likely represent a large portion of the variation contained in the Tumblebrook stock. Our data suggest that genetic drift in these three colonies is actively eroding the standing genetic variation and as they have been maintained in isolation from each other, it has resulted in noticeable differentiation between the colonies.

There are two major ramifications of the loss and partitioning of genetic variation in lab colonies. First, animals from the same original outbred stock may respond very differently to an experiment if they come from different isolated colonies. Many papers state that diversity in gerbils is quite low, one even suggesting that smaller error bars in laboratory individuals than wild-caught individuals are due to the lower genetic diversity (i.e.: Stuermer & Wetzel, 2006). While almost certainly correct that the diversity in their colony is low, our data suggest that is likely a reflection of high drift in an isolated colony, not low diversity in the original Tumblebrook stocks or even across all laboratory gerbils in general. That diversity is low is an important factor in interpreting many experimental results, but generally missing from this acknowledgement is that while diversity is likely low in any specific colony, that does not mean that all colonies are genetically similar. This may partly explain why some experimental outcomes are not able to be replicated despite using animals from the same original outbred strain (Justice & Dhillon, 2016; Richter et al., 2011). This general argument is applicable not only to rodent colonies, but any laboratory animals of any taxa where the population size is limited.

The second important ramification of high genetic drift in laboratory colonies is that while diversity will be lost in any single colony through time, across multiple isolated colonies much of the original diversity may be preserved. This is not a new idea and there are major ongoing efforts using multiple inbred strains to capture the range of natural diversity (Churchill et al., 2004; Collaborative Cross Consortium, 2012; Threadgill & Churchill, 2012). By intercrossing between multiple differentiated colonies, researchers can do controlled experiments designed to uncover the genetic architecture of complex traits (Festing, 2010). In this regard gerbils seem to be ideal candidates. Indeed, the high overall variation, number of private alleles within colonies, and intermediate location of the Sheffield x Edinburgh F1 offspring in the PCA (Figure 2) all suggest that sufficient diversity exists between the colonies for successful genetic experiments.

### Conclusions

Despite being derived from a relatively small number of founders and experiencing repeated bottlenecks over the past 80 years in captivity, the Tumblebrook Farm strain of Mongolian gerbils does not possess low levels of genetic diversity. Genetic drift in small institutional colonies of this species can increase differentiation, and may impact on the reproducibility of results. We advise that experimenters consider the history of their colony when planning and performing research projects using gerbils. We further suggest that published claims on levels of genetic diversity in laboratory rodents based on small numbers of genetic markers should be taken with a pinch of salt.

## ACKNOWLDEGEMENTS

We would like to thank Leila Abbas for providing us with the Edinburgh and Sheffield strains of gerbils and Charlie Parady (Charles River Ltd) and Judith Allen for information on the maintenance of the Charles River and Edinburg gerbil colonies respectively. We thank Matt Hegarty (Aberystwyth University) for advice on GBS sequencing and Rhys Morgan and Emlyn Roberts for technical assistance with animal husbandry. Finally, we would like to thank Jeff Good, Jesse Weber, and Kyle Turner for helping us with diversity estimates from various rodent colonies. This work was funded by a Leverhulme Trust research project grant to J.F.M. and K.A.S. (RPG-2015-450).

## REFERENCES

Abbas, L., & Rivolta, M. N. (2015). Aminoglycoside ototoxicity and hair cell ablation in the adult gerbil: A simple model to study hair cell loss and regeneration. Hearing Research, 325(C), 12–26. http://doi.org/10.1016/j.heares.2015.03.002

Brekke, T. D., & Good, J. M. (2014). Parent-of-origin growth effects and the evolution of hybrid inviability in dwarf hamsters. Evolution, 68(11), 3134–3148. http://doi.org/10.1111/evo.12500

Brekke, T. D., Henry, L. A., & Good, J. M. (2016). Genomic imprinting, disrupted placental expression, and speciation. Evolution, 70(12), 2690–2703. http://doi.org/10.1111/evo.13085

Buchhalter, J. R. (1993). Animal models of inherited epilepsy. Epilepsia, 34(Suppl. 3), S31–S41.

Buckmaster, P. S. (2006). Inherited Epilepsy in Mongolian Gerbils. In A. Pitkänen, P. A. Schwartzkroin, & S. L. Moshé (Eds.), Models of Seizures and Epilepsy (pp. 273–294). Burlington: Academic Press.

Casellas, J. (2011). Inbred mouse strains and genetic stability: a review. Animal, 5(1), 1–7. http://doi.org/10.1017/S1751731110001667

Catchen, J. M., Amores, A., Hohenlohe, P., Cresko, W., & Postlethwait, J. H. (2011). Stacks: building and Genotyping Loci De Novo From Short-Read Sequences. G3: Genes| Genomes| Genetics, 1(3), 171–182. http://doi.org/10.1534/g3.111.000240

Catchen, J., Hohenlohe, P. A., Bassham, S., Amores, A., & Cresko, W. A. (2013). Stacks: an analysis tool set for population genomics. Molecular Ecology, 22(11), 3124–3140.

Chen, W., Jongkamonwiwat, N., Abbas, L., Eshtan, S. J., Johnson, S. L., Kuhn, S., et al. (2012). Restoration of auditory evoked responses by human ES-cell-derived otic progenitors. Nature, 490(7419), 278–282. http://doi.org/10.1038/nature11415

Chia, R., Achilli, F., Festing, M. F. W., & Fisher, E. M. C. (2005). The origins and uses of mouse outbred stocks. Nature Genetics, 37(11), 1181–1186. http://doi.org/10.1038/ng1665

Churchill, G. A., Airey, D. C., Allayee, H., Angel, J. M., Attie, A. D., Beatty, J., et al. (2004). The Collaborative Cross, a community resource for the genetic analysis of complex traits. Nature Genetics, 36(11), 1133–1137. http://doi.org/10.1038/ng1104-1133

Clark, M. M., Ham, M., & Galef, B. G. (1994). Differences in the sex ratios of offspring originating in the right and left ovaries of Mongolian gerbils (*Meriones unguiculatus*). Journal of Reproduction and Fertility, 101(2), 393–396. http://doi.org/10.1016/B978-012263951-7/50010-7

Collaborative Cross Consortium. (2012). The genome architecture of the Collaborative Cross mouse genetic reference population. Genetics, 190, 389–401.

de Koning, D.-J., & McIntyre, L. M. (2017). Back to the Future: Multiparent Populations Provide the Key to Unlocking the Genetic Basis of Complex Traits. Genetics, 206(2), 527–529. http://doi.org/10.1534/genetics.117.203265

Du, X. Y., Li, W., Sa, X. Y., Li, C. L., Lu, J., Wang, Y. Z., & Chen, Z. W. (2015). Selection of an effective microsatellite marker system for genetic control and analysis of gerbil populations in China. Genetics and Molecular Research, 14(3), 11030–11042. http://doi.org/10.4238/2015.September.21.16

Du, X., Chen, Z., Li, W., Tan, Y., Lu, J., Zhu, X., et al. (2010). Development of Novel Microsatellite DNA Markers by Cross-Amplification and Analysis of Genetic Variation in Gerbils. Journal of Heredity, 101(6), 710–716. http://doi.org/10.1093/jhered/esq066

Elshire, R. J., Glaubitz, J. C., Sun, Q., Poland, J. A., Kawamoto, K., Buckler, E. S., & Mitchell, S. E. (2011). A robust, simple genotyping-by-sequencing (GBS) approach for high diversity species. Plos One, 6(5), e19379–e19379. http://doi.org/10.1371/journal.pone.0019379

Eppig, J. T. (2007). Mouse strain and genetic nomenclature: an abbreviated guide. In J. G. Fox, M. T. Davisson, F. W. Quimby, S. W. Barthold, C. E. Newcomer, & A. L. Smith (Eds.), The Mouse in Biomedical Research (2nd ed., Vol. 1, pp. 79–98). London: the mouse in biomedical research.

Falush, D., Stephens, M., & Pritchard, J. K. (2003). Inference of population structure using multilocus genotype data: linked loci and correlated allele frequencies. Genetics, 164(4), 1567–1587.

Falush, D., Stephens, M., & Pritchard, J. K. (2007). Inference of population structure using multilocus genotype data: dominant markers and null alleles. Molecular Ecology Notes, 7(4), 574–578. http://doi.org/10.1111/j.1471-8286.2007.01758.x

Festing, M. F. (1999). Warning: the use of heterogeneous mice may seriously damage your research. Neurobiology of Aging, 20(2), 237–44– discussion 245–6.

Festing, M. F. W. (2010). Inbred Strains Should Replace Outbred Stocks in Toxicology, Safety Testing, and Drug Development. Toxicologic Pathology, 38(5), 681–690. http://doi.org/10.1177/0192623310373776

Fuchsberger, C., Flannick, J., Teslovich, T. M., Mahajan, A., Agarwala, V., Gaulton, K. J., et al. (2016). The genetic architecture of type 2 diabetes. Nature, 536(7614), 1–29. http://doi.org/10.1038/nature18642

Gawriluk, T. R., Simkin, J., Thompson, K. L., Biswas, S. K., Clare-Salzler, Z., Kimani, J. M., et al. (2016). Comparative analysis of ear-hole closure identifies epimorphic regeneration as a discrete trait in mammals. Nature Communications, 7, 1–16. http://doi.org/10.1038/ncomms11164

Groen, A., & Lagerwerf, A. J. (1979). Genic heterogeneity and genetic monitoring of mouse outbred stocks. Laboratory Animals, 13(2), 81–85.

Harper, J. M. (2008). Wild-derived mouse stocks: an underappreciated tool for aging research. Age, 30(2-3), 135–145. http://doi.org/10.1007/s11357-008-9057-0

Justice, M. J., & Dhillon, P. (2016). Using the mouse to model human disease: increasing validity and reproducibility. Disease Models & Mechanisms, 9(2), 101–103. http://doi.org/10.1242/dmm.024547

Li, H., & Durbin, R. (2009). Fast and accurate short read alignment with Burrows-Wheeler transform. Bioinformatics, 25(14), 1754–1760.

Li, H., Handsaker, B., Wysoker, A., Fennell, T., Ruan, J., Homer, N., et al. (2009). The Sequence Alignment/Map format and SAMtools. Bioinformatics, 25(16), 2078–2079.

Li, X., Lu, J., Wang, Y., Huo, X., Li, Z., Zhang, S., et al. (2016). Establishment and Characterization of a Newly Established Diabetic Gerbil Line. Plos One, 11(7), e0159420–13. http://doi.org/10.1371/journal.pone.0159420

McManus, J. J. (1972). Water relations and food consumption of the mongolian gerbil, *Meriones unguiculatus*. Comparative Biochemistry and Physiology Part a: Physiology, 43(4), 959–967. http://doi.org/10.1016/0300-9629(72)90168-5

Meisler, M. H., Kearney, J., Ottman, R., & Escayg, A. (2001). Identification of Epilepsy Genes in Human and Mouse. Annual Review of Genetics, 35(1), 567–588. http://doi.org/10.1146/annurev.genet.35.102401.091142

Ness, R. W., Zhang, Y.-H., Cong, L., Wang, Y., Zhang, J.-X., & Keightley, P. D. (2012). Nuclear Gene Variation in Wild Brown Rats. G3: Genes| Genomes| Genetics 2(12), 1661–1664.

Neumann, K., Maak, S., & Stuermer, I. W. (2001). Low microsatellite variation in laboratory gerbils. Journal of Heredity, 327–347. http://doi.org/10.1002/9781444318777.ch23

Papaioannou, V. E., & Festing, M. F. (1980). Genetic drift in a stock of laboratory mice. Laboratory Animals, 14(1), 11–13. http://doi.org/10.1258/002367780780943015

Petrij, F., van Veen, K., Mettler, M., & Brückmann, V. (2001). A second acromelanistic allelomorph at the albino locus of the Mongolian gerbil (Meriones unguiculatus). Journal of Heredity, 92(1), 74–78.

Pritchard, J. K., Stephens, M., & Donnelly, P. (2000). Inference of population structure using multilocus genotype data. Genetics, 155(2), 945–959.

Razzoli, M., Papa, R., Valsecchi, P., & Nonnis Marzano, F. (2003). AFLP to Assess Genetic Variation in Laboratory Gerbils (*Meriones unguiculatus*). Journal of Heredity, 94(6), 507–511.

Rich, S. S. (2016). Diabetes: Still a geneticist’s nightmare. Nature, 536(7614), 37–38. http://doi.org/10.1038/nature18906

Richter, S. H., Garner, J. P., Zipser, B., Lewejohann, L., Sachser, N., Touma, C., et al. (2011). Effect of Population Heterogenization on the Reproducibility of Mouse Behavior: A MultiLaboratory Study. Plos One, 6(1), e16461–14. http://doi.org/10.1371/journal.pone.0016461

Rosenberg, N. A. (2003). distruct: a program for the graphical display of population structure. Molecular Ecology Notes, 4(1), 137–138. http://doi.org/10.1046/j.1471-8286.2003.00566.x

Rosenberg, N. A., Pritchard, J. K., Weber, J. L., Cann, H. M., Kidd, K. K., Zhivotovsky, L. A., & Feldman, M. W. (2002). Genetic Structure of Human Populations. Science (New York, N.Y.) 298(5602), 2381–2385. http://doi.org/10.2307/3833180?ref=search-gateway:7b5294a290ca39cb8bef6104439e20fb

Russell, W. M. S., & Burch, R. L. (1959). The Principles of Humane Experimental Technique. London: Methuen & Co. Ltd.

Salcedo, T., Geraldes, A., & Nachman, M. W. (2007). Nucleotide Variation in Wild and Inbred Mice. Genetics 177(4), 2277–2291.

Smits, B. M. G., van Zutphen, B. F. M., Plasterk, R. H. A., & Cuppen, E. (2004). Genetic variation in coding regions between and within commonly used inbred rat strains. Genome Research, 14(7), 1285–1290.

Srivastava, A., Morgan, A. P., Najarian, M. L., Sarsani, V. K., Sigmon, J. S., Shorter, J. R., et al. (2017). Genomes of the Mouse Collaborative Cross. Genetics, 206(2), 537–556. http://doi.org/10.1534/genetics.116.198838

Stuermer, I. W., & Wetzel, W. (2006). Early experience and domestication affect auditory discrimination learning, open field behaviour and brain size in wild Mongolian gerbils and domesticated Laboratory gerbils (*Meriones unguiculatus forma domestica*). Behavioural Brain Research, 173(1), 11–21.

Stuermer, I. W., Plotz, K., Leybold, A., Zinke, O., Kalberlah, O., Samjaa, R., & Scheich, H. (2003). Intraspecific Allometric comparison of Laboratory gerbils with Mongolian Gerbils Trapped in the Wild Indicates Domestication in *Meriones unguiculatus* (Milne-Edwards, 1867) (Rodentia: Gerbillinae). Zoologischer Anzeiger - a Journal of Comparative Zoology, 242(3), 249–266. http://doi.org/10.1078/0044-5231-00102

Svenson, K. L., Gatti, D. M., Valdar, W., & Welsh, C. E. (2012). High-resolution genetic mapping using the Mouse Diversity outbred population. Journal of Heredity. http://doi.org/10.1534/genetics.111.132597/-/DC1

Thiessen, D. D., & Kittrell, E. M. (1980). The Harderian gland and thermoregulation in the gerbil (*Meriones unguiculatus*). Physiology & Behavior, 24(3), 417–424. http://doi.org/10.1016/0031-9384(80)90229-2

Threadgill, D. W., & Churchill, G. A. (2012). Ten years of the collaborative cross. Genetics. http://doi.Org/10.1534/genetics.111.138032/-/DC1/FileS1.pdf

Vincent, A. L., & Rodrick, G. E. (1979). The pathology of the Mongolian Gerbil (*Meriones unguiculatus*): a review. Laboratory Animal Science, 29(5), 645–651.

Wang, D., Wang, Y., & Wang, Z. (2000). Metabolism and thermoregulation in the Mongolian gerbil *Meriones unguiculatus*. Acta Theriologica, 45(2), 183–192. http://doi.org/10.4098/AT.arch.00-21

Weber, J. N., Peterson, B. K., & Hoekstra, H. E. (2013). Discrete genetic modules are responsible for complex burrow evolution in Peromyscus mice. Nature, 493(7432), 402–405. http://doi.org/10.1038/nature11816

Zheng, X., Levine, D., Shen, J., & Gogarten, S. M. (2012). A high-performance computing toolset for relatedness and principal component analysis of SNP data. Journal of Heredity, 28(24), 3326–3328.

